# Molecular epidemiological study of Scrub Typhus in residence, farm and forest habitats from Yunnan Province, China

**DOI:** 10.1101/2023.07.13.548801

**Authors:** Jia-Wei Tian, Yi-Chen Kong, Pei-Yu Han, Fen-Hui Xu, Wei-Hong Yang, Yun-Zhi Zhang

## Abstract

The number of people suffering from scrub typhus, which is not of concern, is increasing year by year, especially in Yunnan Province, China. From June 1, 2021 to August 15, 2022, a total of 505 mammalian samples were collected from farm, forest, and residential habitats with high incidence of scrub typhus in Yunnan, China, for nPCR (nested PCR) and qPCR (quantitative real-time PCR) detection of *Orientia tsutsugamushi*. A total of 4 orders of murine-like animals, Rodentia (87.52%, n=442), Insectivora (10.29%, n=52), Lagomorpha (1.79%, n=9) and Scandentia (0.40%, n=2) were trapped. Comparing the qPCR infection rates in the three habitats, it was no significant difference that the infection rate of residential habitat (44.44%) and that of the farm habitat (45.05%, P>0.05), which is much larger than that of the forest habitat (3.08%) (P<0.001). Three genotypes (Karp, Kato and TA763) of *O. tsutsugamushi* were found from Yunnan, China in this study. We found a tendency for scrub typhus to migrate from forests to residential habitats.

**Author Summary:** Scrub typhus is acute febrile infectious disease caused by *Orientia tsutsugamushi* that threatens nearly 1 billion people. According to the data we have obtained, the incidence rate of infected people has reached 23.84/100,000 population until December 2022 in Yunnan, China. The incidence rate has increased non-negligibly Compared with 0.65/100,000 population in 2006. Among them, the incidence rate has increased by 71.14% from 2019 to 2022. Scrub typhus is transmitted by the bite of chigger larvae with murine-like animals as the main source of infection Rodentia are the most important reservoir hosts, followed by Insectivora. Therefore, in view of the influencing factor of human habitat, we used qPCR (quantitative real-time PCR) and nPCR (nested PCR) techniques to analyze the prevalence of *O. tsutsugamushi* in hosts of different human habitats from June 2021 to August 2022. Our research shows that the diversity of *O. tsutsugamushi* genotypes in Yunnan Province provides guidance for the prevention and Control of scrub typhus. And it was found that the infection rate of *O. tsutsugamushi* in murine-like animals is quite different in different human habitats, especially the high infection rate in residential habitat and farm habitat, suggesting that *O. tsutsugamushi* can be infected without wilderness history.

## 1. Introduction

As a re-emerging unclassified acute febrile infectious disease, scrub typhus, caused by *Orientia tsutsugamushi*, has a serious impact on the public health of Asia-Pacific countries. The World Health Organization has declared scrub typhus one of the most underdiagnosed/underreported diseases in the world. It has been stated two decades ago that more than 1 million cases of scrub typhus occur each year and 1 billion people are at risk of disease exposure in an habitat of more than 13 million square kilometers endemic for scrub typhus [1]. In southern China, the predicted high-risk habitats for scrub typhus transmission are mainly concentrated in the five Province of Yunnan, Guangxi, Guangdong, Hainan and Fujian. It is estimated that 162.684 million people live in potential infection risk habitats in southern China [2]. From 2006 to 2016, the total number of reported and treated cases of scrub typhus in China increased from 254 to 21,562, and the annual incidence rate also increased dramatically, from 0.09/100,000 to 1.60/100,000 population [3]. In Yunnan province, which accounts for nearly a quarter of all scrub typhus infections in China, clinical diagnosis mostly relies on field activity history, as sensitive laboratory diagnosis of scrub typhus is difficult in resource-limited habitats, especially in border regions [4]. It is believed that a better understanding of the epidemic situation will help guide policymakers to formulate effective regional control strategies. This study conducted a host risk assessment of scrub typhus to determine the topography and direction of disease transmission in Yunnan Province, China.

The genus *Orientia* belongs to the order Rickettsiales within the family Rickettsiaceae. Chigger mites are currently the only known vector of scrub typhus. They have four life cycles: eggs, larvae (often called chiggers), nymphs, and adults. Except for the parasitic life of larvae, they live in the soil for the rest of the period. However, *O. tsutsugamushi* is vectored by the biting of the larval life stage of infected chigger mites (e.g. *Leptotrombidium deliense, L*.*scutellare*) [5]. The species diversity of chigger mites in Yunnan was much higher than diversities reported previously in the other Provinces of China and in other countries. Small mammals (e.g. rodents and insectivores) are the most common hosts of chiggers [6]. They play an important role in the transmission of scrub typhus in nature as they can transport scrub typhus-carrying chiggers from endemic to non-endemic habitats. Scrub typhus is often transmitted to humans through the bite of chigger mites carried by small mammals [7]. Therefore, the impact of scrub typhus on humans can be judged and the risk estimated by studying the infection rate of small mammals.

## 2. Methods

### 2.1 Data collection and sampling

We obtained the number of scrub typhus cases in Yunnan Province from 2006 to 2022 through the Yunnan Institute of Endemic Diseases Control and Prevention, and we obtained population data for calculating incidence from the National Bureau of Statistics of China (http://data.stats.gov.cnenglish/index.htm). We calculated annual incidence using the number of human scrub typhus cases divided by the corresponding population number at the end of a given year.

Using fried food as bait and placing mouse traps, the small mammal investigation was carried out from farm, forest and residential habitats in Yunnan Province, China from June 1, 2021 to August 15, 2022 (Fig. 1). The collected animal samples were immediately brought back to the laboratory for morphological identification. Rodents were euthanized and dissected to collect the heart, liver, spleen, lung, kidney, and intestinal tissues placed in 2 mL cryogenic vials (CORNING, China) which were stored at -80°C for further analysis. The mitochondrial cytochrome b (mt-Cytb) gene of liver tissue DNA was amplified by PCR for molecular biological identification [8]. The species, geographical location, altitude and air humidity of the captured animals were recorded.

**Figure 1.**
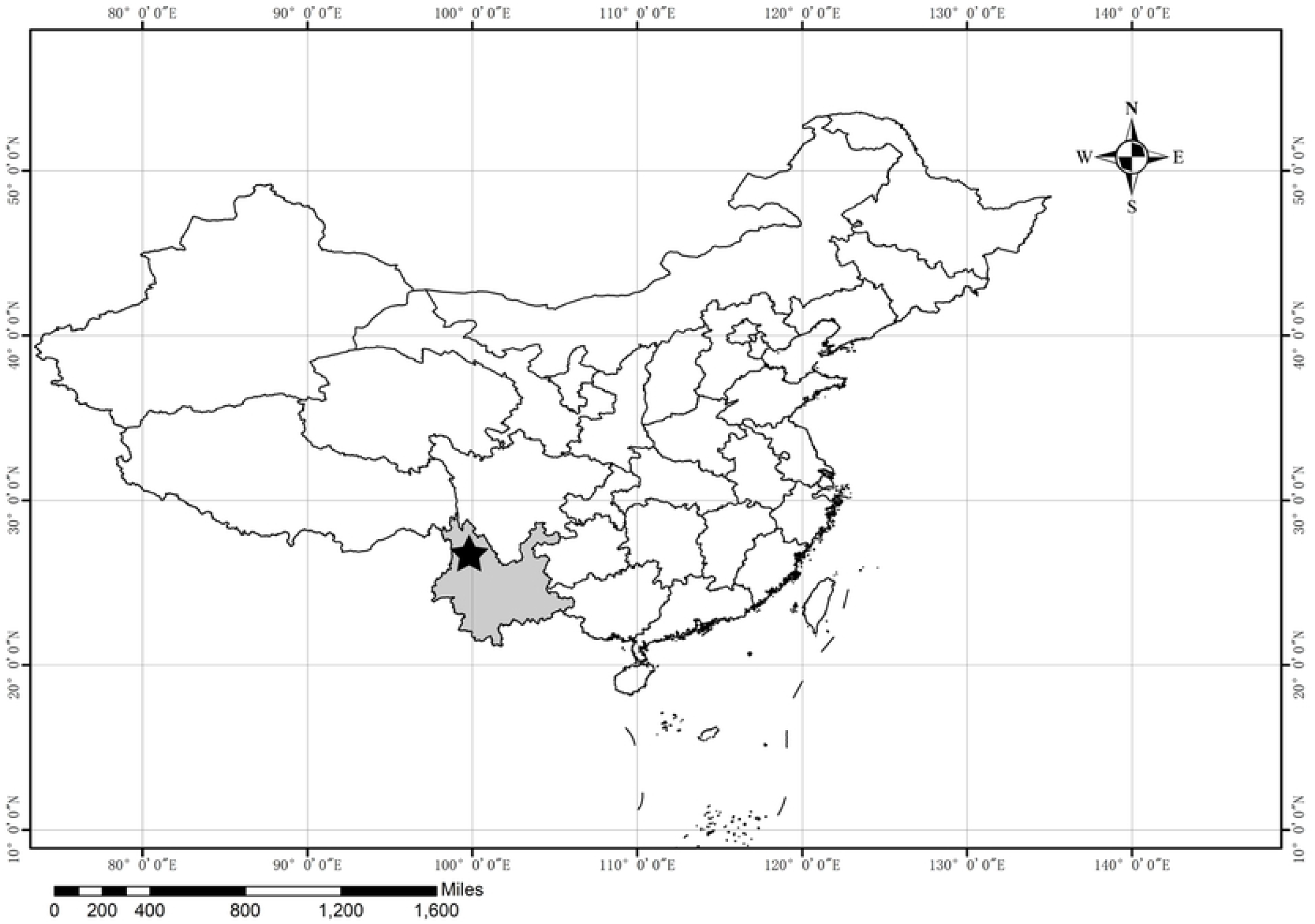
*Orientia tsutsugamushi* in small mammals, Yunnan Province, China, June 2021-August 2022. The asterisk represents the location for this study of Yunnan, China.

### 2.2 Nucleic acid extraction

The collected mammalian spleen tissue for detection were aseptically clipped, and about 0.1 g of spleen tissue was placed into the GeneReady animal PIII pulverization tube (Suizhen, China) to which sterilized 500 μL of 7.4 phosphate-buffered saline (PBS) had been added. Grinding was performed in a GeneReady Ultimate biological sample cryogenic rapid preparation centrifuge system (Suizhen, China). Nucleic acid extraction from animal tissue grinding fluid (200μL) was performed using a DNA extraction kit (TIANGEN, China) according to the manufacturer’s instructions.

### 2.3 Specimens detection

The genotype of *O. tsutsugamushi* in host animals was determined by nested polymerase chain reaction (nPCR) using the 56kDa outer membrane protein genome [9].Two sets of primers used were as follows: outer primers, Ot-Out-F1

(5’-TACATTAGCTGCGGGTATGACA-3’), Ot-OutR1

(5’-CCAGCATAATTCTTCAACCAAG-3’) and inner primers Ot-In-F2

(5’-GAGCAGAGCTAGGTGTTATGTA-3’), Ot-In-R2

(5’-TAGGCATTATAGTAGGCTGAGG-3’).

The 47kDa gene which has amplified 1401bp product was amplified using genomic DNA of the *O. tsutsugamushi* Karp strain as the template. *E. coli* cells containing target DNA on LB medium were grown in LB medium for 9 hours. Plasmid DNA was extracted from 600μl of the suspension, using the Plasmid Mini Kit I (Omega Bio-Tek, America), following the manufacturer’s instructions and the concentration was determined. The *O. tsutsugamushi* data in ng/μl were then converted to numbers of copies/μl. For plasmid detection sensitivity, we performed real-time quantitative polymerase chain reaction (qPCR) with serial dilutions of plasmid DNA from 5×10^9^ copies/μl to 5×10^3^ copies/μl using DNase-Free Deionized Water to create a standard curve.

The presence of *O. tsutsugamushi* was assessed by qPCR and the degree of infection in the host animals was determined using the 47kDa high temperature protein genome. qPCR was accomplished using probe (FAM-TGGGTAGCTTTGGTGGACCGATGTTTA ATCT-BHQ1) to determine *O. tsutsugamushi* copy numbers [10].

### 2.4 Phylogenetic and data analysis

All data were statistically analyzed using SPSS 24.0 statistical software. The Chi-square test was used for statistical methods, and P <0.05 was considered statistically significant.

The measured sequence was compared with the nucleotide sequence of *O. tsutsugamushi* in GenBank by BLAST algorithm and the nucleotide and amino acid sequences of similar strains were obtained. The sequence was compared with 7 similar strains using the BioAider program [11] (Additional file: Table S1), and then the phylogenetic tree was constructed using the neighbor-joining method (NJ method) in the MEGA `X` software. The bootstrap value was 1000, and the distance was determined by the maximum likelihood method calculate.

## 3. Results

The total number of scrub typhus cases in Yunnan Province increased from 292 to 11,189, and the annual incidence rate increased from 0.65/100,000 to 23.84/100,000 population from 2006 to 2022. As of December 2022, compared to the end of 2019, the incidence rate of infections has grown by 71.14% from 13.93/100,000 to 23.84/100,000 population (Fig. 2).

**Figure 2.**
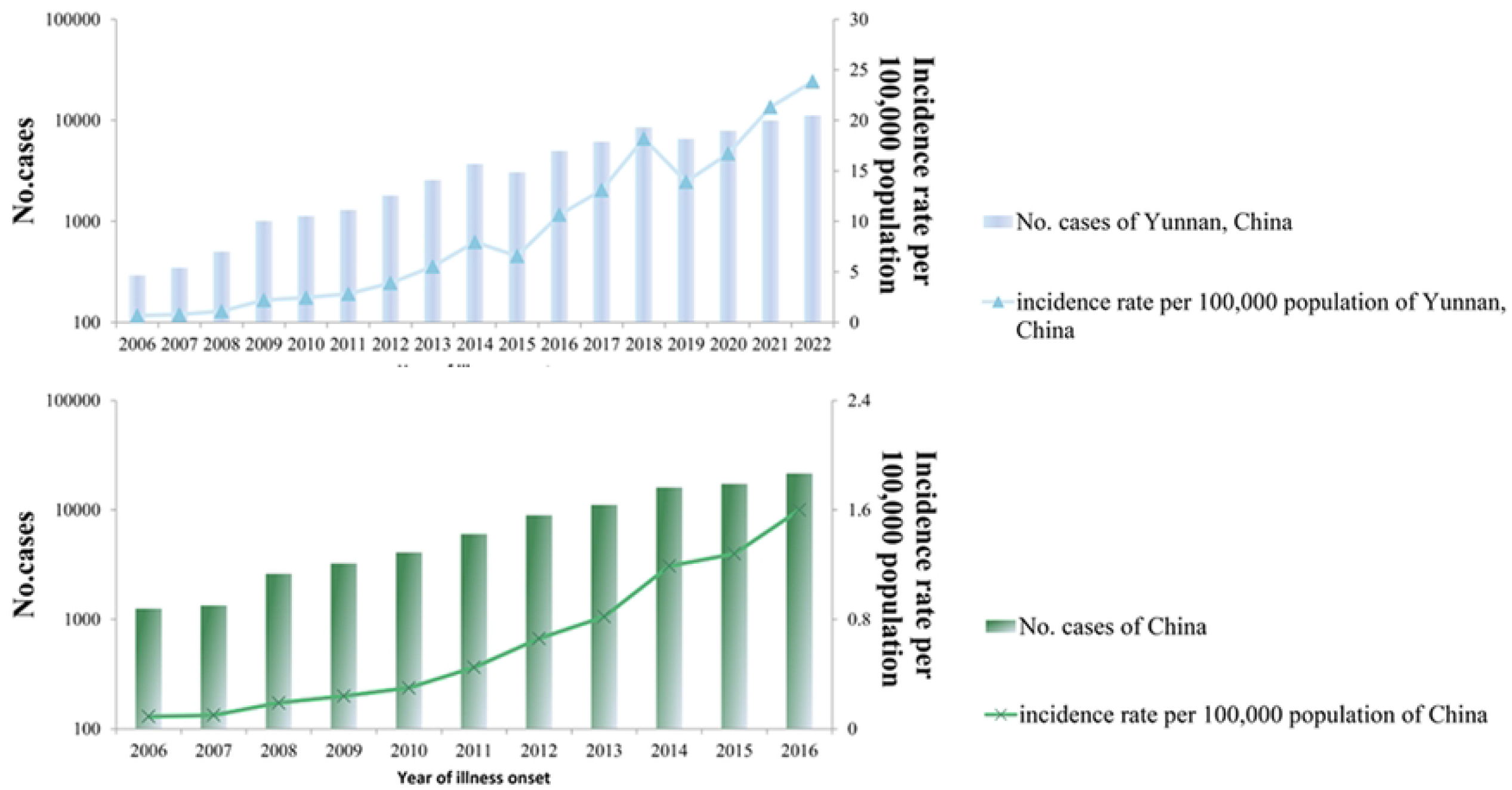
Reported cases of scrub typhus. (A) Cases in China, 2006–2016. (B) Cases in Yunnan, China, 2006–2022.

A total of 505 mammal samples were collected from farm habitat (36.04%, n=182), forest habitat (51.49%, n=260) and residential habitat (12.47%, n=63) which were collected from high-incidence habitats of scrub typhus in Yunnan Province, China from June 1, 2021 to August 15, 2022.

We captured a total of 4 orders of murine-like animals: Rodentia (87.52%, n=442), Insectivora (10.29%, n=52), Lagomorpha (1.79%, n=9) and Scandentia (0.40%, n=2). The dominant species in the farm, residential and forest habitats are the Chevrieri’s field mouse (*Apodemus chevrieri*), Asian house rat (*Rattus tanezumi*) and *A. chevrieri*, respectively. High infection rates in murine-like animals were observed in qPCR results in residential and farm habitats. Comparing the qPCR infection rates in the three habitats, it was no significant difference that the infection rate of residential habitat (44.44%) and that of the farm habitat (45.05%) (P>0.05), which are much larger than that of the forest habitat (3.08%) (P<0.001). The total positive rate of qPCR was 23.56%.

The nPCR infection rate in the residential habitat was consistent with that in the farm habitat, both of which were higher than those in the forest habitat (P<0.001) (Table 1). The nucleotide and amino acid sequences of the 56kDa gene sequences obtained in this study were compared with similar strains (Additional file: Table S2).

**Table 1.**
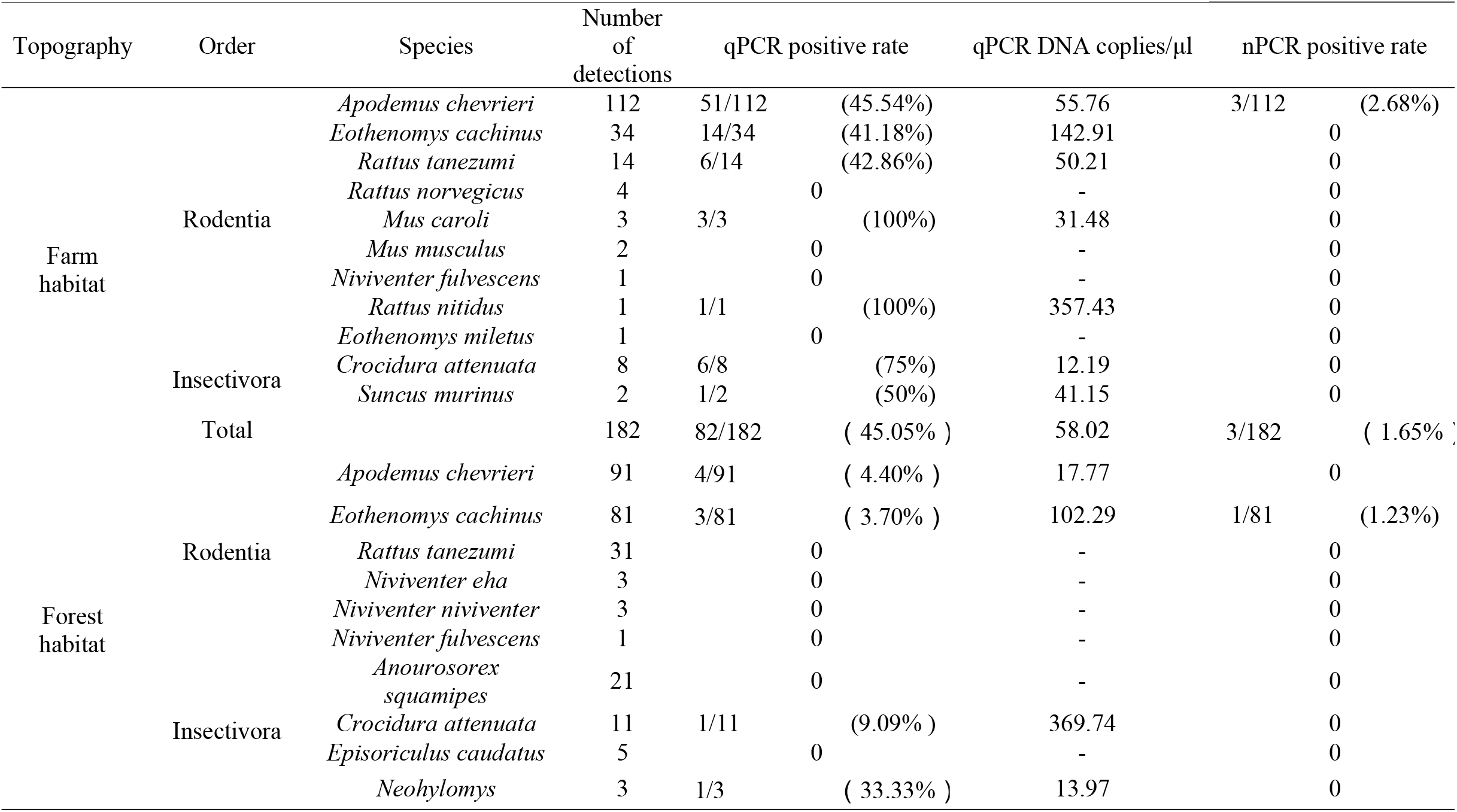
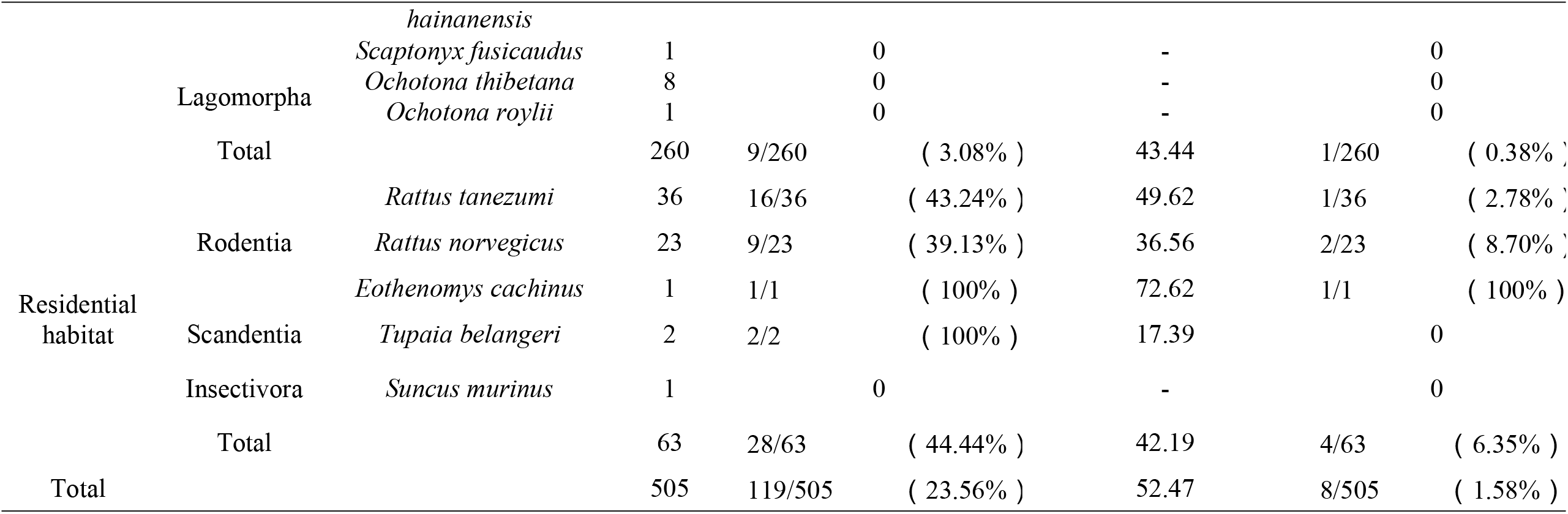
Prevalence of *Orientia tsutsugamushi* in small mammals, Yunnan Province, China, June 2021-August 2022

Phylogenetic analysis showed that the three (Accession no: OP925103-OP925105) and four (OM914542, OP925100-OP925102) of the sequences were clustered with Kato genotypes, TA763 genotypes respectively and one sequence (OP925009) clustered with Karp genotype identified by 56kDa in this study (Fig. 3). A total of three genotypes were detected in this study. The nucleotide homology between a Karp genotype *O. tsutsugamushi* detected in this experiment and a patient from Yunnan reached 98.8%. The three detected TA763 strains had 98.8%-99.4% nucleotide homology with the patient and chigger from Taiwan. In addition, there are three strains of Kato genotype scrub typhus, which share 98.65%-99.32% nucleotide homology with patients from Yunnan and Thailand respectively.

**Figure 3.**
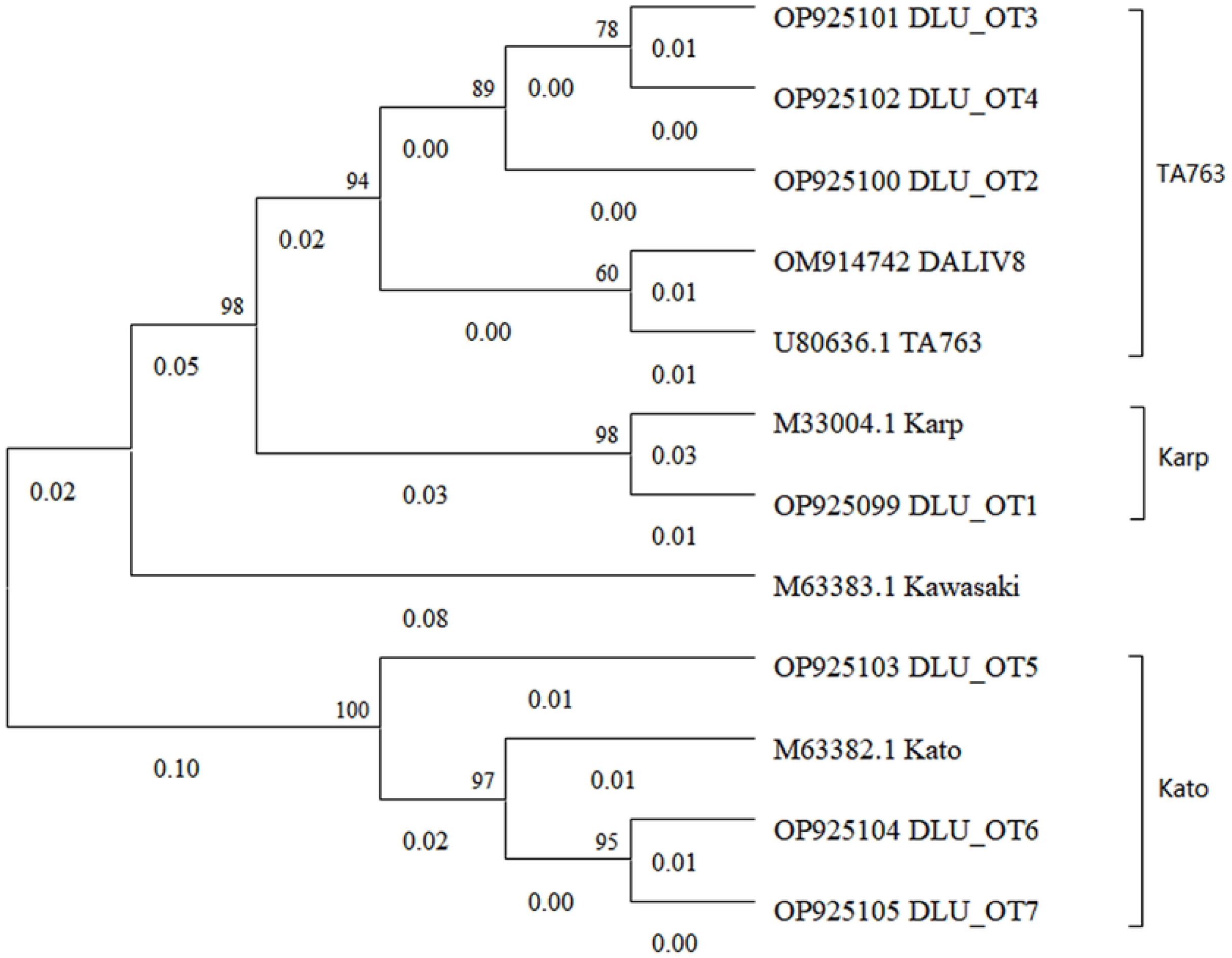
The phylogenetic tree of this study.

The infection rate of *O. tsutsugamushi* in murine-like animals was positively correlated with the disturbance level of human activities, contrary to the findings of previous years. However, this result is consistent with the increase in the number of infections in Yunnan over the same period.

## 4. Discussion

This study collected the latest scrub typhus case data, and found that both the number of scrub typhus cases and the incidence rate are still on the rise by the end of 2022 in Yunnan, China. This requires us to identify the potential relationship between scrub typhus and its related factors as soon as possible, so as to curb the rapid spread of this disease, which poses a major threat to human health. Therefore, we collected murine-like animal samples to explore the relationship between scrub typhus and human habitats.

Among the 505 samples in this experiment, three genotypes were found to reflect the diversity of *O. tsutsugamushi* genotypes in Yunnan, China. The TA763 genotype was first isolated from *Rattus rajah* in Thailand [12]. The TA763 sequence obtained in this study is the most similar to the sequence obtained in Taiwan which can be considered that *O. tsutsugamushi* may have been migrated over long distances. In addition, there are three strains of Kato genotype, which are most similar to patients in Yunnan and Thailand respectively. However, researchers have investigated that the most common genotypes are Karp and Karp-related in Thailand [13, 14]. In the reports from China, the dominant genotypes of scrub typhus in Yunnan were Karp and Kato. Different genotypes of *O. tsutsugamushi* have a tendency to spread within “tsutsugamushi triangle”[15]. This result suggests that 56kDa sequence detection plays a role in establishing cross-border emerging infectious disease linkages. In this experiment, the dominant genotypes of scrub typhus are considered as Kato and TA763 in Yunnan Province during 2020-2022.

The terrain of Yunnan is dominated by plateaus [16]. This experiment found that the infection rate in forest habitats is lower than that in farm land and residential habitats. It may be considered that the forests in Southwest China are dominated by high altitudes, and the lower temperature in high altitude habitats is not suitable for the survival of chiggers [17, 18]. In addition, Yunnan Province is adjacent to several countries with high incidence of scrub typhus [19-21]. Therefore, it is necessary to study scrub typhus, an emerging infectious disease, because the incidence rate in Yunnan Province has been increasing year by year. In a previous study of hosts and parasites in Yunnan Province, the researchers collected 10,222 small mammals of 62 species that were infested by 92,990 chiggers of 224 species [22]which can observe the species diversity of scrub typhus hosts and vectors in Yunnan Province. Emerging zoonotic infectious diseases such as scrub typhus may be the main mode of transmission of infectious diseases in the future. In animal hosts, emerging pathogens may not cause symptoms in host animals. However, while these pathogens are not always in the host animal, the pathogen evolves in small animals until it spills over into humans [23]. Scrub typhus is mainly transmitted to humans through the bites of pathogenic chiggers parasitic on small mammals, after contacting humans through small mammals [24]. Our results indicate that the infection rate of rodents is higher in habitats with high human activity than in habitats with low human activity, and the year-on-year increase in the incidence of scrub typhus in Yunnan may be due to the influence of in such varying environments.

We found an increased risk of exposure to *O. tsutsugamushi* in residential habitats. The difference of infection rate of *O. tsutsugamushi* in hosts indicated that scrub typhus tended to move from forests to farm and into cities. In 2021, an ecological analysis of scrub typhus in Thailand found that, forest habitat was poorly associated with *O. tsutsugamushi* infection in all the analyses [18]. This finding is inconsistent with the cognition of previous articles on the discovery of scrub typhus infection [25]. This may explain one reason of the increasing number of scrub typhus cases in Yunnan Province year by year [26]. The reason for this is speculated to be the reduction of forest habitat in Yunnan and the replacement of forests with farm or residential habitats [27]. The second reason is to suspect that the emergence of global warming is causing geographic migration of rodent hosts carrying scrub typhus [28]. The occurrence of both of these situations will have the result of the migration of rodents to residential habitats [29]. It is suggested that the risk of scrub typhus may change significantly under the rapid environmental changes in humans.

## 5. Conclusions

We found a tendency for scrub typhus to migrate from forests to residential habitats.

## Declarations

## Acknowledgments

We would like to thank Yunnan Institute of Endemic Diseases Control and Prevention for their help in collecting samples and Data for this study.

## Authors’ contributions

J.W.T and Y.Z.Z contributed to the study design. Y.C.K and P.Y.H collected the data. F.H.X and W.H.Y analyzed the data, which was interpreted by all authors. J.W.T wrote the manuscript. Y.Z.Z gave some useful comments and suggestions to this work. All authors reviewed the manuscript. All authors approved the final manuscript.

## Ethics approval and consent to participate

This research was approved by the Medical Ethics Committee of Dali University under number DLDXLL2018008 and was obtained with the informed consent of all participants.

## Consent for publication

Not applicable.

## Author details

Mr. Tian is a master student at Dali University, mainly engaged in the research of vector-borne diseases.

## References

1. Watt G, Parola P. Scrub typhus and tropical rickettsioses. Curr Opin Infect Dis. 2003;16:429–436. https://doi.org/10.1097/00001432-200310000-00009 PMID: 14501995

2. Zheng C, Jiang D, Ding F, Fu J, Hao M. Spatiotemporal Patterns and Risk Factors for Scrub Typhus From 2007 to 2017 in Southern China. Clin Infect Dis. 2019;69:1205–1211. https://doi.org/10.1093/cid/ciy1050 PMID: 30535175

3. Li Z, Xin H, Sun J, Lai S, Zeng L, Zheng C, et al. Epidemiologic Changes of Scrub Typhus in China, 1952-2016. Emerg Infect Dis. 2020;26:1091–1101. https://doi.org/10.3201/eid2606.191168 PMID: 32441637

4. Saraswati K, Day NPJ, Mukaka M, Blacksell SD. Scrub typhus point-of-care testing: A systematic review and meta-analysis. PLoS Negl Trop Dis. 2018;12:e0006330. https://doi.org/10.1371/journal.pntd.0006330 PMID: 29579046

5. Mullen GR, Durden LA. Medical and veterinary entomology. Academic press; 2009.

6. Alkathiry H, Al-Rofaai A, Ya’cob Z, Cutmore TS, Mohd-Azami SNI, Husin NA, et al. Habitat and Season Drive Chigger Mite Diversity and Abundance on Small Mammals in Peninsular Malaysia. Pathogens. 2022;11:1087. https://doi.org/10.3390/pathogens11101087 PMID: 36297144

7. Kuo CC, Lee PL, Chen CH, Wang HC. Surveillance of potential hosts and vectors of scrub typhus in Taiwan. Parasit Vectors. 2015;8:611. https://doi.org/10.1186/s13071-015-1221-7 PMID: 26626287

8. Guillen-Servent A, Francis CM. A new species of bat of the Hipposideros bicolor group (Chiroptera : Hipposideridae) from Central Laos, with evidence of convergent evolution with Sundaic taxa. Acta Chiropterologica. 2006;8:39–61. https://doi.org/10.3161/1733-5329(2006)8[39:Ansobo]2.0.Co;2.

9. Furuya Y, Yoshida Y, Katayama T, Yamamoto S, Kawamura A, Jr. Serotype-specific amplification of Rickettsia tsutsugamushi DNA by nested polymerase chain reaction. J Clin Microbiol. 1993;31:1637–1640. https://doi.org/10.1128/JCM.31.6.1637-1640.1993 PMID: 8315007

10. Jiang J, Chan TC, Temenak JJ, Dasch GA, Ching WM, Richards AL. Development of a quantitative real-time polymerase chain reaction assay specific for Orientia tsutsugamushi. Am J Trop Med Hyg. 2004;70:351–356. https://doi.org/10.4269/ajtmh.2004.70.351 PMID: 15100446

11. Zhou ZJ, Qiu Y, Pu Y, Huang X, Ge XY. BioAider: An efficient tool for viral genome analysis and its application in tracing SARS-CoV-2 transmission. Sustain Cities Soc. 2020;63:102466. https://doi.org/10.1016/j.scs.2020.102466 PMID: 32904401

12. Phuklia W, Panyanivong P, Sengdetka D, Sonthayanon P, Newton PN, Paris DH, et al. Novel high-throughput screening method using quantitative PCR to determine the antimicrobial susceptibility of Orientia tsutsugamushi clinical isolates. J Antimicrob Chemother. 2019;74:74–81. https://doi.org/10.1093/jac/dky402 PMID: 30295746

13. Blacksell SD, Luksameetanasan R, Kalambaheti T, Aukkanit N, Paris DH, McGready R, et al. Genetic typing of the 56-kDa type-specific antigen gene of contemporary Orientia tsutsugamushi isolates causing human scrub typhus at two sites in north-eastern and western Thailand. FEMS Immunol Med Microbiol. 2008;52:335–342. https://doi.org/10.1111/j.1574-695X.2007.00375.x PMID: 18312580

14. Ruang-Areerate T, Jeamwattanalert P, Rodkvamtook W, Richards AL, Sunyakumthorn P, Gaywee J. Genotype diversity and distribution of Orientia tsutsugamushi causing scrub typhus in Thailand. J Clin Microbiol. 2011;49:2584–2589. https://doi.org/10.1128/jcm.00355-11 PMID: 21593255

15. Richards AL, Jiang J. Scrub Typhus: Historic Perspective and Current Status of the Worldwide Presence of Orientia Species. Trop Med Infect Dis. 2020;5:49. https://doi.org/10.3390/tropicalmed5020049 PMID: 32244598

16. Han H, Liang Y, Song Z, He Z, Duan R, Chen Y, et al. Epidemiological Characteristics of Human and Animal Plague in Yunnan Province, China, 1950 to 2020. Microbiol Spectr. 2022;10:e0166222. https://doi.org/10.1128/spectrum.01662-22 PMID: 36219109

17. Steventon C, Harley D, Wicker L, Legione AR, Devlin JM, Hufschmid J. An assessment of ectoparasites across highland and lowland populations of Leadbeater’s possum (Gymnobelideus leadbeateri): Implications for genetic rescue translocations. Int J Parasitol Parasites Wildl. 2022;18:152–156. https://doi.org/10.1016/j.ijppaw.2022.05.002 PMID: 35586791

18. Elliott I, Thangnimitchok N, Chaisiri K, Wangrangsimakul T, Jaiboon P, Day NPJ, et al. Orientia tsutsugamushi dynamics in vectors and hosts: ecology and risk factors for foci of scrub typhus transmission in northern Thailand. Parasit Vectors. 2021;14:540. https://doi.org/10.1186/s13071-021-05042-4 PMID: 34663445

19. Elders PND, Dhawan S, Tanganuchitcharnchai A, Phommasone K, Chansamouth V, Day NPJ, et al. Diagnostic accuracy of an in-house Scrub Typhus enzyme linked immunoassay for the detection of IgM and IgG antibodies in Laos. PLoS Negl Trop Dis. 2020;14:e0008858. https://doi.org/10.1371/journal.pntd.0008858 PMID: 33284807

20. Elders PND, Swe MMM, Phyo AP, McLean ARD, Lin HN, Soe K, et al. Serological evidence indicates widespread distribution of rickettsioses in Myanmar. Int J Infect Dis. 2021;103:494–501. https://doi.org/10.1016/j.ijid.2020.12.013 PMID: 33310022

21. Trung NV, Hoi LT, Dien VM, Huong DT, Hoa TM, Lien VN, et al. Clinical Manifestations and Molecular Diagnosis of Scrub Typhus and Murine Typhus, Vietnam, 2015-2017. Emerg Infect Dis. 2019;25:633–641. https://doi.org/10.3201/eid2504.180691 PMID: 30882318

22. Zhan YZ, Guo XG, Speakman JR, Zuo XH, Wu D, Wang QH, et al. Abundances and host relationships of chigger mites in Yunnan Province, China. Med Vet Entomol. 2013;27:194–202. https://doi.org/10.1111/j.1365-2915.2012.01053.x PMID: 23167491

23. Olival KJ, Hosseini PR, Zambrana-Torrelio C, Ross N, Bogich TL, Daszak P. Host and viral traits predict zoonotic spillover from mammals. Nature. 2017;546:646–650. https://doi.org/10.1038/nature22975 PMID: 28636590

24. Meerburg BG, Singleton GR, Kijlstra A. Rodent-borne diseases and their risks for public health. Crit Rev Microbiol. 2009;35:221–270. https://doi.org/10.1080/10408410902989837 PMID: 19548807

25. Lin EC, Tu HP, Hong CH. Halved Incidence of Scrub Typhus after Travel Restrictions to Confine a Surge of COVID-19 in Taiwan. Pathogens. 2021;10:1386. https://doi.org/10.3390/pathogens10111386 PMID: 34832542

26. Peng PY, Xu L, Wang GX, He WY, Yan TL, Guo XG. Epidemiological characteristics and spatiotemporal patterns of scrub typhus in Yunnan Province from 2006 to 2017. Sci Rep. 2022;12:2985. https://doi.org/10.1038/s41598-022-07082-x PMID: 35194139

27. Shi L, Dossa GGO, Paudel E, Zang H, Xu J, Harrison RD. Changes in Fungal Communities across a Forest Disturbance Gradient. Appl Environ Microbiol. 2019;85: e00080–19. https://doi.org/10.1128/AEM.00080-19 PMID: 30979833

28. Wei Y, Huang Y, Li X, Ma Y, Tao X, Wu X, et al. Climate variability, animal reservoir and transmission of scrub typhus in Southern China. PLoS Negl Trop Dis. 2017;11:e0005447. https://doi.org/10.1371/journal.pntd.0005447 PMID: 28273079

29. Roberts T, Parker DM, Bulterys PL, Rattanavong S, Elliott I, Phommasone K, et al. A spatio-temporal analysis of scrub typhus and murine typhus in Laos; implications from changing landscapes and climate. PLoS Negl Trop Dis. 2021;15:e0009685. https://doi.org/10.1371/journal.pntd.0009685 PMID: 34432800

